# Heat activation and inactivation of bacterial spores. Is there an overlap?

**DOI:** 10.1101/2021.11.20.469368

**Authors:** Juan Wen, Jan P. P. M. Smelt, Norbert O.E. Vischer, Arend L. de Vos, Peter Setlow, Stanley Brul

**Affiliations:** Molecular Biology and Microbial Food Safety, Swammerdam Institute for Life Sciences, University of Amsterdam, Science Park 904, 1098 XH Amsterdam, The Netherlands; Department of Molecular Biology and Biophysics, UConn Health, 263 Farmington Avenue, Farmington, CT 06030-3305, United States of America

**Keywords:** *Bacillus subtilis*, bacterial spores, heat activation, heat inactivation, spore germination, germination heterogeneity

## Abstract

Heat activation at a sublethal temperature is widely applied to promote *Bacillus* species spore germination. This treatment also has potential to be employed in food processing to eliminate undesired bacterial spores by enhancing their germination, and then inactivating the less heat resistant germinated spores at a milder temperature. However, incorrect heat treatment could also generate heat damage in spores, and lead to more heterogeneous spore germination. Here, the heat activation and heat damage profile of *Bacillus subtilis* spores was determined by testing spore germination and outgrowth at both population and single spore levels. The heat treatments used were 40-80°C, and for 0-300 min. The results were as follows. 1) Heat activation at 40-70°C promoted L-valine and L-asparagine-glucose-fructose-potassium (AGFK) induced germination in a time dependent manner. 2) The optimal heat activation temperatures for AGFK and L-valine germination via the GerB plus GerK or GerA germinant receptors were 65 and 50-65°C, respectively. 3) Heat inactivation of dormant spores appeared at 70°C, and the heat damage of molecules essential for germination and growth began at 70 and 65°C, respectively. 4) Heat treatment at 75°C resulted in both activation of germination and damage to the germination apparatus, and 80°C treatment caused more pronounced heat damage. 5) For the spores that should withstand adverse environmental temperatures in nature, heat activation seems functional for a subsequent optimal germination process, while heat damage affected both germination and outgrowth.

## Introduction

Applying mild heat for a short to moderate time (e.g. pasteurization treatment at 72°C for at least 15 s) is one of the conventional food processing procedures to extend shelf life in the food industry [1,2]. In this process, vegetative pathogens or spoilage organisms lose their viability, although bacterial spores survive. Not only that, heat treatment at sublethal temperatures (eg. 60-75°C) can increase and synchronize the spore germination of *Bacillales* and *Clostridiales* [2–5]. *Bacillus subtilis* spores have higher heat resistance than their vegetative forms due to the following factors [2–4]. 1). The spore core has a low water content (25-45% wet weight) and a high level of dipicolinic acid (DPA) chelated to divalent cations, predominantly Ca^2+^ (Ca^2+^DPA; 25% of core dry weight). 2) Spore DNA is saturated with **α**/**β**-type small acid-soluble protein (SASP). 3) Spore cortex peptidoglycan, characterized by the muramic-δ-lactam moiety and low level of peptide cross-linking, also contributes to spore wet heat resistance. However, upon increasing the temperature above 80°C, spore damage and inactivation occur [6]. Damage accumulated in spore molecules then results in more heterogeneous germination and slowed spore outgrowth [6]. To analyze spore thermal physiology such that heat activation can be optimally applied for the stimulation of spore germination, it is necessary to have a detailed analysis of the effects of heat treatment, even at relatively mild temperatures.

Heat activation studies have mainly focused on spores of *Bacillales* [5,7–11]. These studies have indicated that a sublethal heat treatment can reversibly activate spores’ germinant receptor (GR) dependent germination, which is induced by physiological small molecule nutrients or hydrostatic pressures of ∼150 MPa. In contrast, sublethal thermal treatment has no effect on GR-independent germination triggered by exogenous Ca^2+^DPA, dodecylamine, or hydrostatic pressures of ∼550 MPa [4,9,12]. In the heat activation process, temperature is not the only variable, as treatment time also affects outcomes, as different treatment times result in different outcomes. Luu *et al*. indicated that the best activation of *B. subtilis* spores was reached after 15 minutes at 75°C with L-valine triggered germination via the GerA GR, whereas a 4 hours at 75°C was needed for optimal germination in L-asparagine, glucose, fructose and potassium (AGFK) triggered germination via the GerB and GerK GRs [9]. When both heat treatment temperature and time duration are considered, the following questions need to be addressed. 1). What is the best time-temperature combination for heat activation? 2). When does heat activation shift to heat inactivation during long treatment times?

In the current paper, we employed *B. subtilis* PS832 spores to measure the time– temperature–activation/inactivation profile at sublethal temperatures. Spores were treated at 40-80°C for 15-300 min, followed by inducing germination by L-valine or AGFK. The spore viability after a variety of treatments was also tested to probe for heat inactivation. Spore germination, and subsequent outgrowth and the growth were monitored at the population level by microplate spectrophotometric analysis, and at the single cell level by phase contrast microscopy.

## Results

### 1. Heat treatments at 40-70°C promote *B. subtilis* spore germination in a time dependent manner

Heat activation at sublethal temperatures is a common procedure to promote homogeneous spore germination in the laboratory. However, the applied temperature varies among different labs, with 65, 70, 75 and 80°C used for *B. subtilis* spore heat activation [9,13–15]. In addition, as mentioned above, heat treatments at 75°C enhance AGFK induced spore germination in a time dependent manner [9]. However, it is not clear whether different temperatures promote germination in a similar pattern. Firstly, we focused on the four commonly used heat activation temperatures mentioned above. As shown in (**Fig. 1D-G**), except for 80°C, the other three temperatures promoted AGFK-induced germination in a time dependent positive manner. However, 65°C treatment resulted in the largest decrease in the optical density at 595 nm (OD_595_), which represents the most complete germination. To test the germination promotion efficiency at lower temperatures, the same measurements were taken on spores treated at 40, 50 and 60°C. As shown in (**Fig. 1A-C**), the time positive dependent correlation was observed again. According to the data presented here the optimum condition of heat activation for AGFK induced germination when completion of germination was monitored was 300 min at 65°C. Notably, the magnitude of the drop in the OD_595_ gradually decreased as the temperature was increased, and was completely abolished at 80°C (**Fig. 1G**). These results suggested that the activation of germination by heat might vanish at 80°C, or was subsumed by damage at this temperature. Similar results, albeit with subtle differences, were observed for L-valine induced germination (**Fig. 2A-G**). First, the optimal heat activation temperature for L-valine induced germination was 50-65°C. Second, treatment at 75°C, while clearly promoting L-valine induced germination, also decreased the magnitude of the maximum OD_595_ decrease. These results suggested there is a transition from only heat activation to heat activation plus damage to spores affecting germination around 75°C.

**Figure 1.**
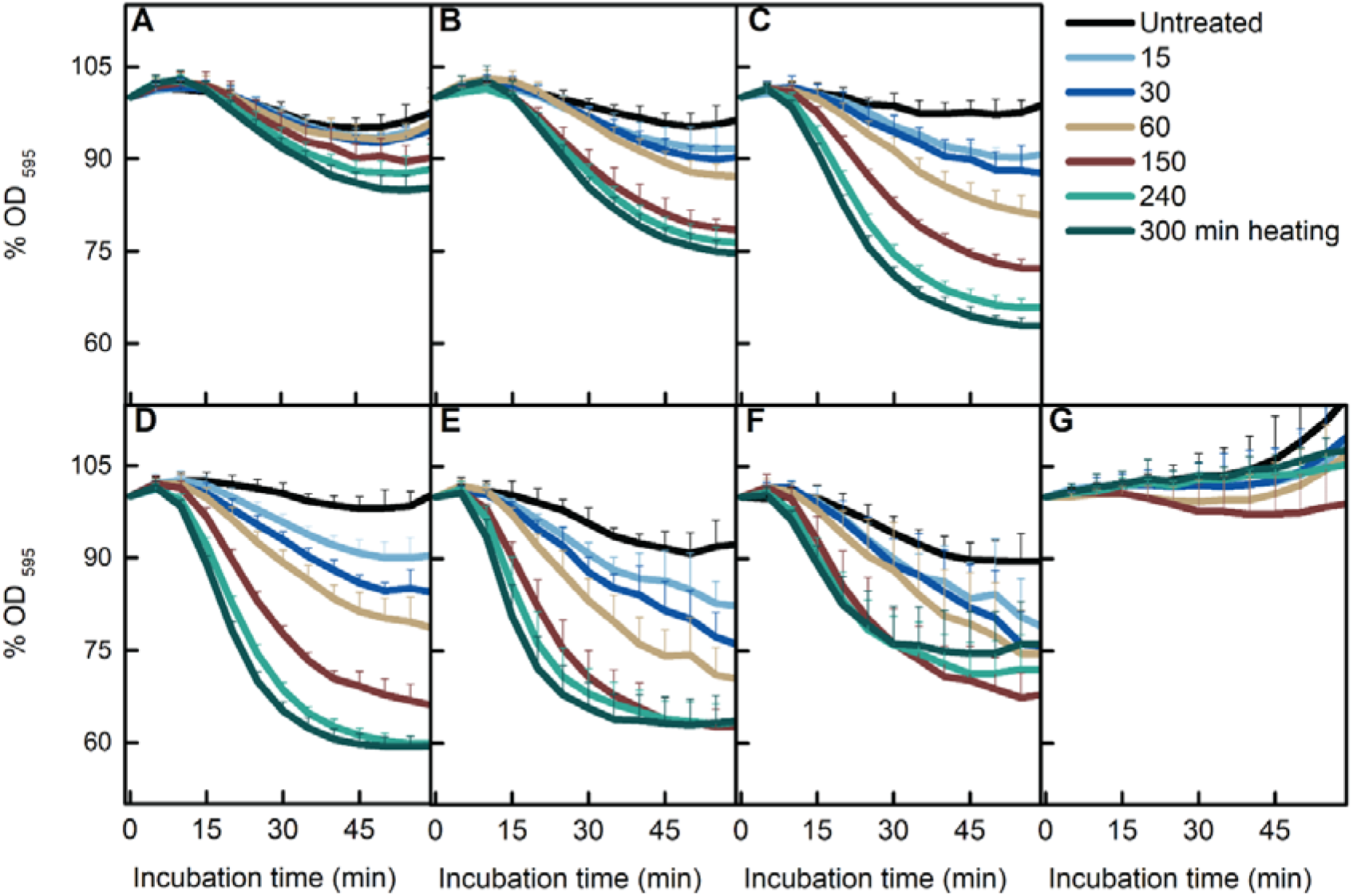
Effect of heat treatment on AGFK induced spore germination. *B. subtilis* PS832 spores were germinated at 37°C with (10 mM each) AGFK in HEPES buffer after heat treatments for various times (0, 15, 30, 60, 150, 240, and 300 min) at 40 (A), 50 (B), 60 (C), 65 (D), 70 (E), 75 (F), and 80°C (G), respectively. Spore germination was measured by the drop in the optical density due to the release of Ca^2+^DPA and water uptake. Values shown are the mean values of duplicate or triplicate measurements on at least two experiments.

**Figure 2.**
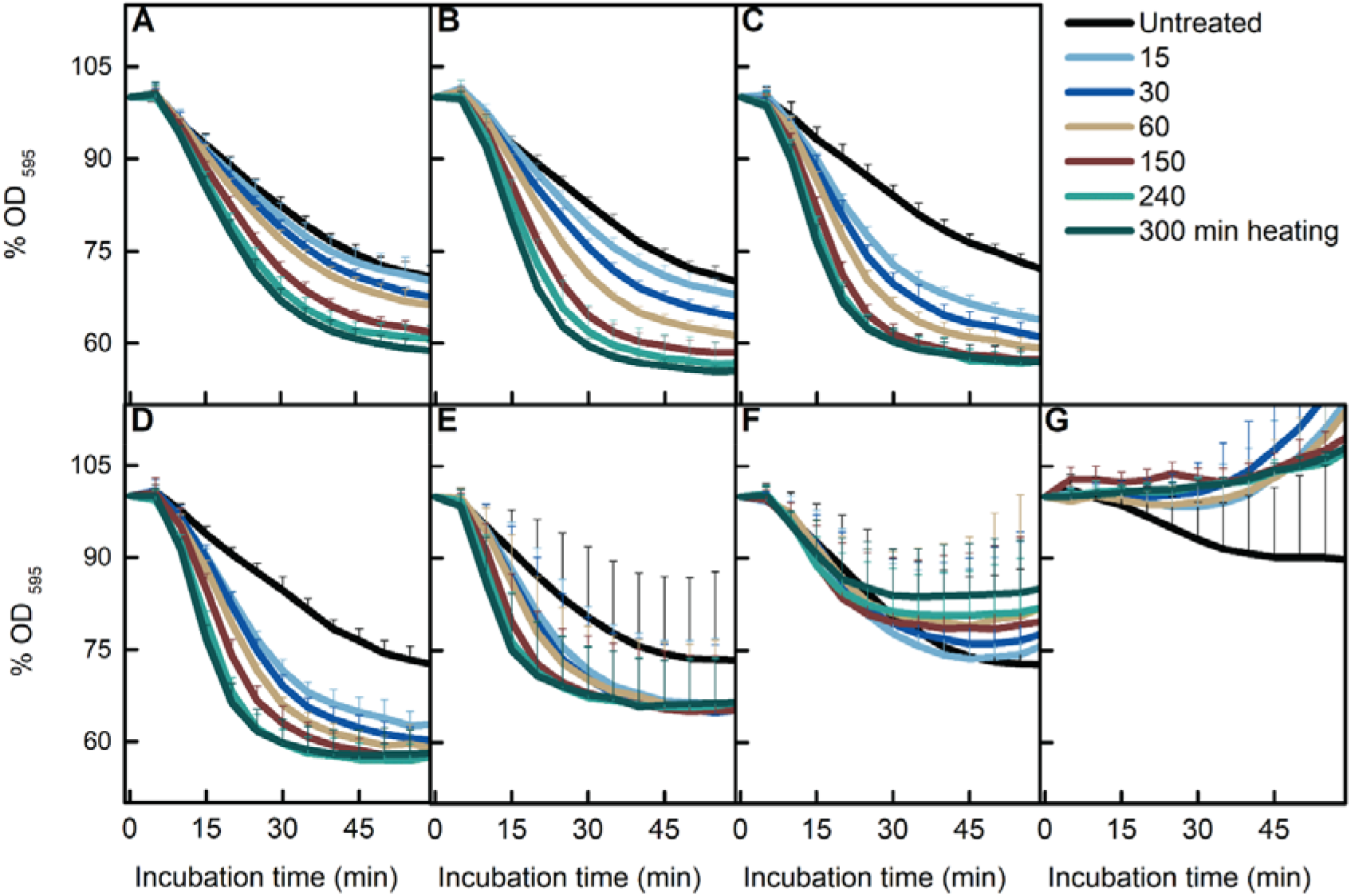
Effect of heat treatment on L-valine induced spore germination. *B. subtilis* PS832 spores were germinated at 37°C with 10 mM L-valine in HEPES buffer after heat treatment for various times (0, 15, 30, 60, 150, 240, and 300 min) at 40 (A), 50 (B), 60 (C), 65 (D), 70 (E), 75 (F), or 80°C (G), respectively. Spore germination was measured by the drop in optical density due to the release of Ca^2+^DPA and water uptake. Values shown are the mean values of duplicate or triplicate measurements in two or three experiments.

### 2. Heat damage accumulates at 70°C

To determine the temperature where heat damage occurs, we tested the viability of spores treated at 65-80°C for various times. Clearly, 80°C treatment led to significant killing of spores (**Fig. 3**). Careful analysis of the results of the other three temperatures, also showed a small, but significant decrease in spore viability for the spores treated for 300 min at 70°C (**Fig. 3B**). To test the effect of heat treatments on cell growth, we tracked the change of spore OD_595_ in LB medium supplemented with AGFK or L-valine. Thermal treatments of 60 and 70°C did not affect *B. subtilis* growth, whereas 75 and 80°C exposure at longer heat treatment times (150, 240, and 300 min) decreased the yield of growing cells when LB medium was supplemented with either L-valine or AGFK (**Fig. 4A-H**). These results suggested that both damage and inactivation of spores take place at 70-75°C.

**Figure 3.**
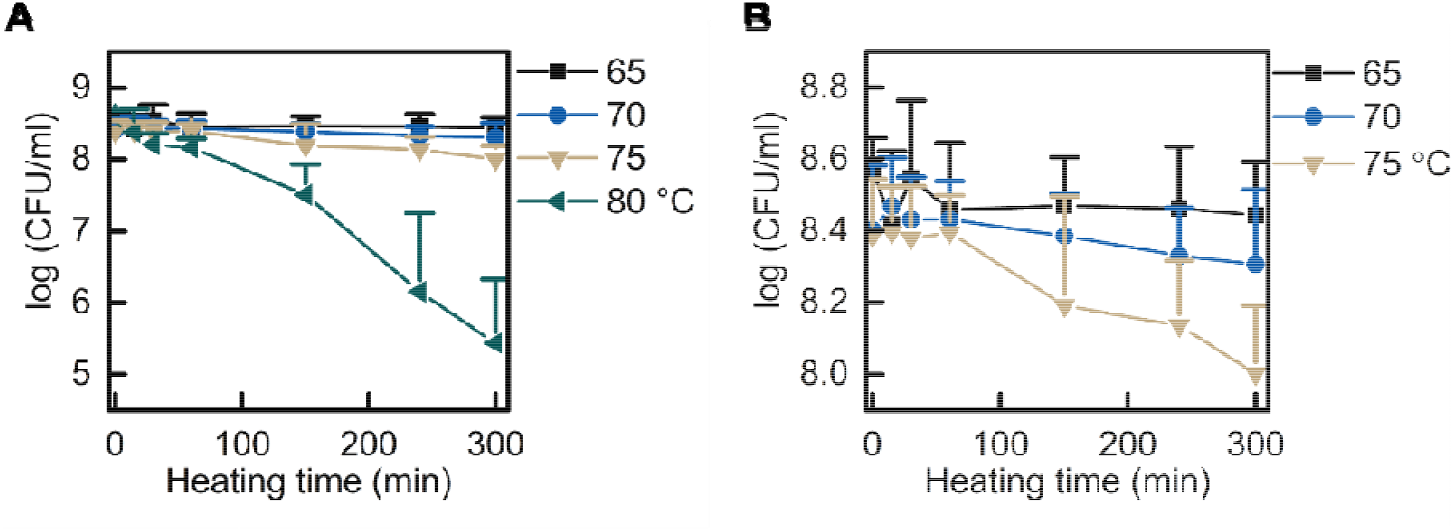
Effect of heat treatment on spore viability. *B. subtilis* PS832 spores were incubated at 37°C on LB agar plates supplemented with (10 mM each) AGFK after heat treatment at 65, 70, 75, or 80°C for various times (0, 15, 30, 60, 150, 240, and 300 min). (A) Viability of spores pre-treated at 65, 70, 75, and 80°C. (B) An enlarged portion of panel A. Values shown are the mean values of triplicate measurements in two or three experiments.

**Figure 4.**
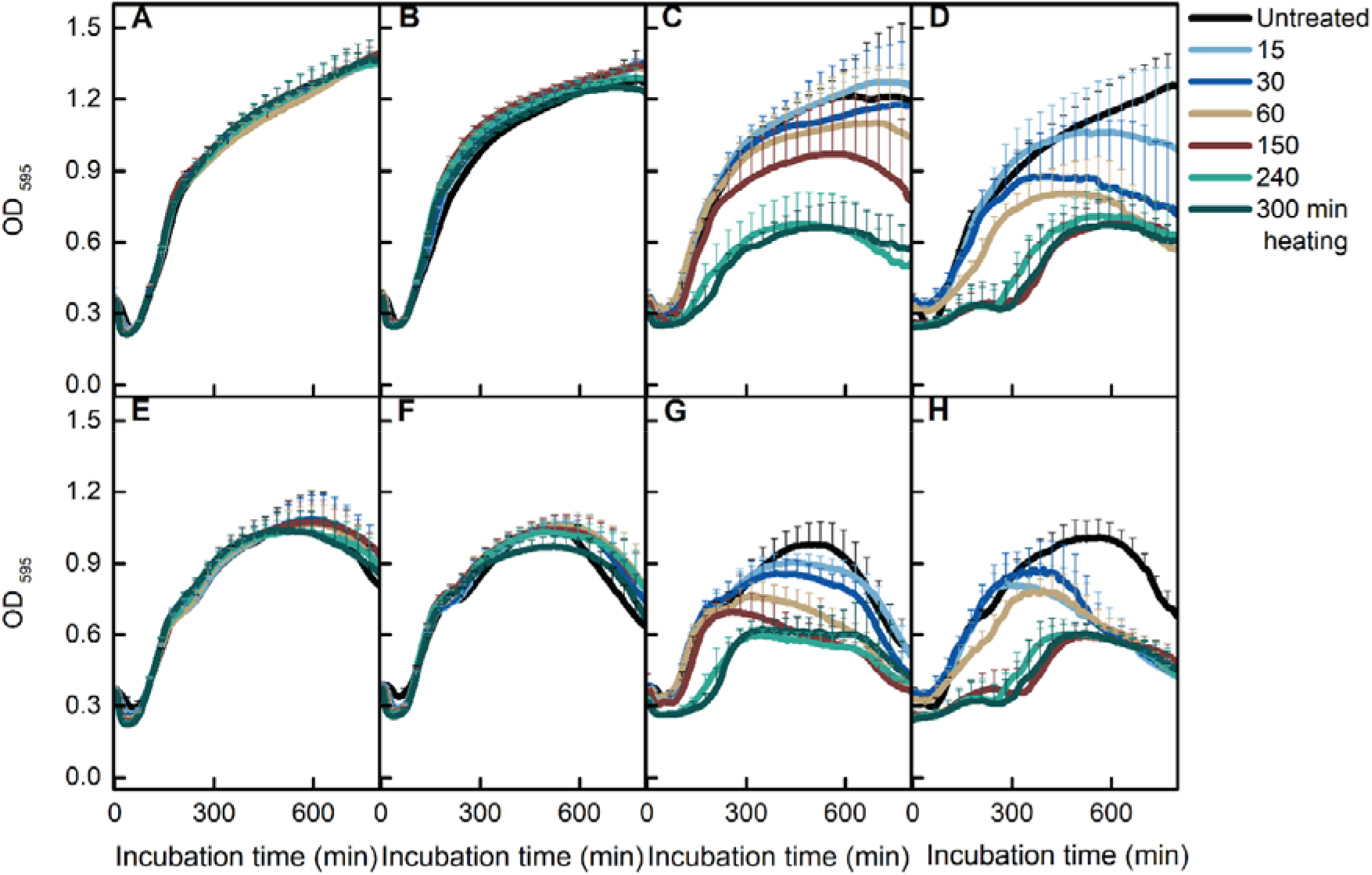
Effect of heat treatment on spore germination and cell growth. *B. subtilis* PS832 spores were heated for various times (0, 15, 30, 60, 150, 240, and 300 min) at 65 (A and E), 70 (B and F), 75 (C and G), and 80°C (D and H), respectively. Spore germination and cell growth were monitored by the change of optical density in LB medium supplemented with (10 mM each) AGFK (A-D) or 10 mM L-valine (E-H). Values shown are the means of triplicate measurements in two or three experiments.

### 3. Effects of heat treatment at single spore level

Our population data indicated that treatments at 65-75°C resulted in a time positive enhancement of AGFK induced spore germination at the population level, and detectable heat damage/inactivation were observed in spores treated at 70-75°C for long times. Considering that the population level data cannot provide a detailed view of what sublethal heat does to individual *B. subtilis* spores, we employed phase contrast microscopy to track AGFK induced spore germination, outgrowth, and cell growth after treatments at 65, 70, 75, and 80°C. By analyzing the time lapse images, the germination plots and growth plots of individual spores were created by SporeTrackerX (ImageJ Macro) (**Fig. 5**). Based on the change of plot profiles, germination, burst (the cell escape from the spore coat), and first cell division events were detected and marked for further analysis. Heat treatment durations at 65-70°C were 30, 150, and 300 min, while treatments for 15 and 60 min at 80°C were added, because of a lack of sufficient germination events for quantitative analysis in the groups with prolonged heat treatment (**Table 1**).

**Figure 5.**
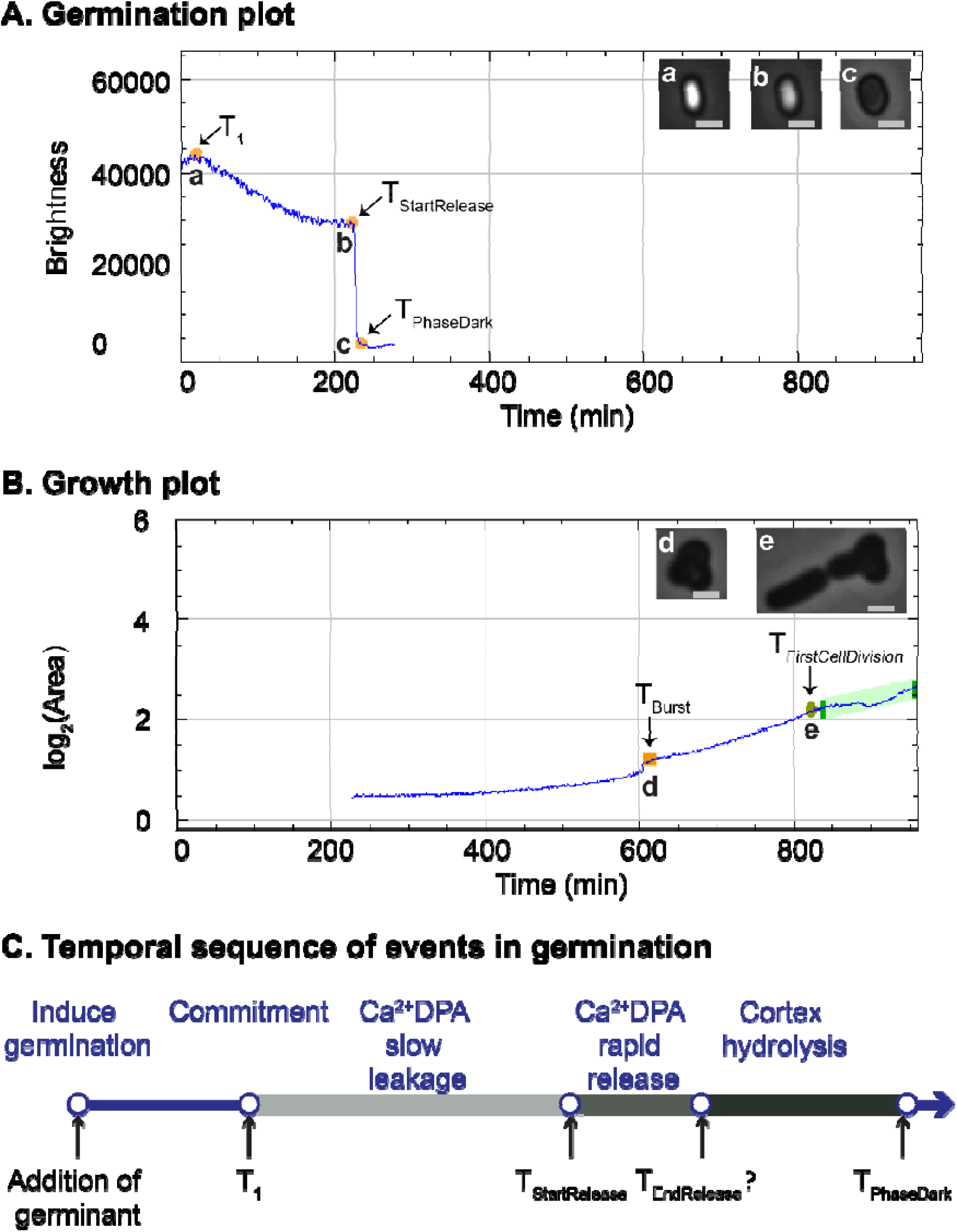
Germination and growth plot of a single *B. subtilis* spore created by SporeTrackerX. In the germination plot (A), two phases of brightness drop at different speeds were observed. The slow drop was considered as the slow leakage of Ca^2+^DPA. The fast drop was likely due to both the rapid Ca^2+^DPA release and the subsequent cortex hydrolysis induced by the released Ca^2+^DPA. The time of the start of rapid Ca^2+^DPA release (T_StartRelease_), time of complete phase darkening (T_PhaseDark_), as well as the time of the start of the slow drop in brightness (T_1_) were assessed during spore germination by SporeTrackerX. The time of burst (T_Burst_) and time of first cell division (T_FirstCellDivision_) were determined, and the various events were marked in the growth plot (B). Micrographs of spores and cells at each time point are shown at the upper-right of A and B. The brightness display range for images a-b and c-e are 0-40000 and 0-25000, respectively. Scale bar, 1 µm. (C). The temporal sequence of events in germination, are adapted from the work of Wang *et al* [19]. The upper part shows the temporal sequence of germination events, and lower part indicates the time points detected in current work as shown in panel A. For the rapid Ca^2+^DPA release, we only estimated its initiation from the germination plot.

**Table 1.**
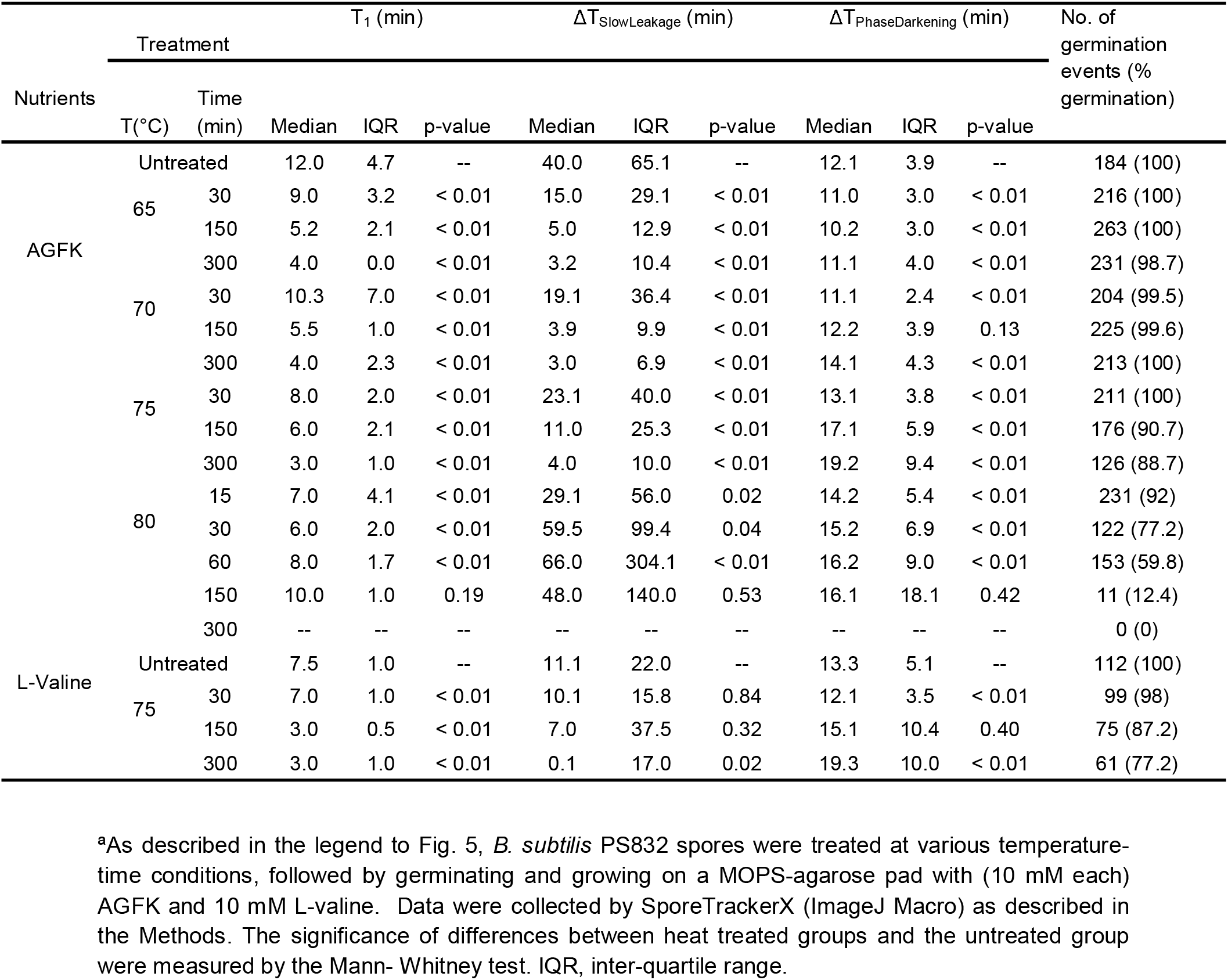
Germination parameters of individual heat treated *B. subtilis* PS832 spores in time lapse images^a^.

#### Damage to the germination apparatus occurs at 70°C

As shown in **Fig. 5A**, the spore germination plot can be divided into three segments: before brightness drop, followed by a slow drop in brightness, and a subsequent fast drop in brightness. Wang *et al*. observed slow leakage of Ca^2+^DPA in *B. subtilis* spore germination, along with the slow decrease of differential interference contrast image intensity, followed by the rapid release of remaining Ca^2+^DPA and cortex hydrolysis during spore germination, and they further suggested that spores become committed to germinate at the starting point of the slow Ca^2+^DPA leakage [17]. We speculated that the slow decrease of spore brightness in the germination plot is due to the slow leakage of Ca^2+^DPA, and T_1_ is where spore commitment occurs (**Fig. 5C**). The fast brightness drop in our germination plot was attributed to the rapid release of Ca^2+^DPA and cortex hydrolysis triggered by release Ca^2+^DPA. Thus, we defined the corresponding duration for each segment in the germination plot as T_1_, ΔT_SlowLeakage_, ΔT_PhaseDarkening_, respectively. Our live imaging data showed that AGFK induced germination was shortened by heat treatments at 65-75°C in a time positive dependent manner (**Fig. 6A-D**). The T_PhaseDark_ of untreated spores, and those treated at 65-75°C for 300 min was 66.6, 21.1, 22.1, and 30.2 min, respectively. In addition, 65-75°C treatments shortened T_1_ and ΔT_SlowLeakage_ in a time positive manner, and 70-75°C treatments prolonged the ΔT_PhaseDarkening_ in a time positive manner (**Table 1**). Given the fact that the release of Ca^2+^DPA is via the SpoVA channel in the spore inner membrane and is slowed markedly when the cortex lytic enzyme CwlJ is absent, SpoVA proteins and cortex lytic enzymes are potential heat damage targets when the thermal treatment temperatures rise above 70°C [16].

**Figure 6.**
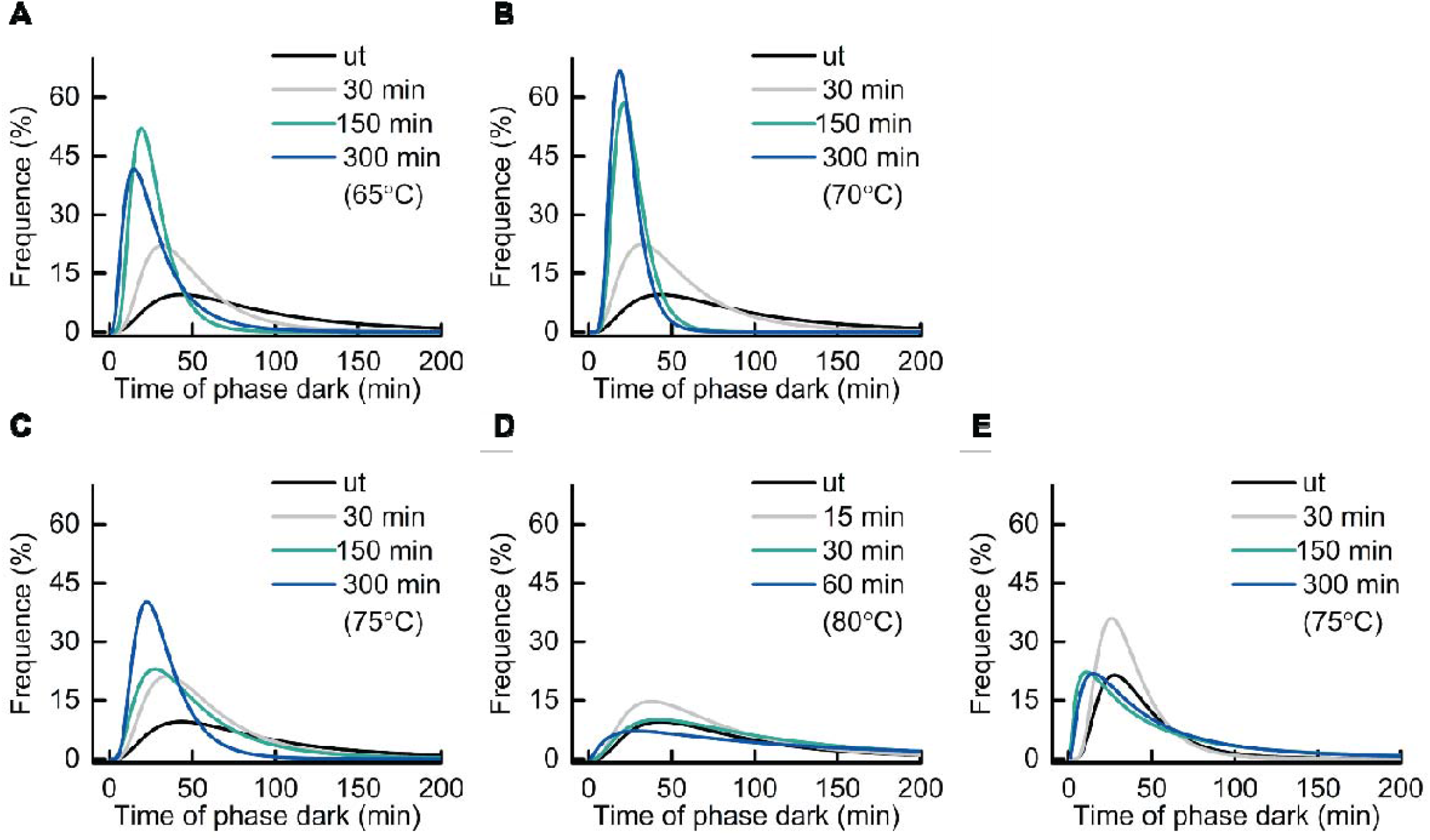
Frequency distribution of the time of the appearence of phase dark spore(T_PhaseDark_). *B. subtilis* PS832 spores were preheated at 65, 70, 75, or 80°C for various times, or not heated (ut). Subsequently, spores were immobilized on a MOPS-Agarose pad at 37°C with (10 mM each) AGFK (A-D) or 10 mM L-valine (E) for 16 hours of time lapse imaging. The line profile is the log-normal distribution fit of the time to phase darkening. The number of germinated spores examined by microscopy are given in Table 1.

#### 65°C treatment can be sufficient to damage essential cell growth molecules stored in spores

The accumulated heat damage, potentially in the germination apparatus, is likely responsible for the T_PhaseDark_ distribution of spores treated at 75 and 80°C (**Fig. 6C, D**). In addition, a decreased germination efficiency was observed in spores treated at 75 and 80°C, and this decline has a time/temperature positive correlation (**Table 1, Fig. 7**). The subsequent outgrowth and cell growth of heat-treated spores also has been monitored in our time lapse imaging. We noticed a remarkable decrease (∼41%) of the cell division events of spores treated at 70°C for 300min compared to untreated spores, although 100% of spores of both groups germinated (**Table 1, 2, Fig. 7**). The decline of cell division events occurred in spores treated at 65°C for 150 min, and increased when the treatment temperature and time increased (**Table 1, 2, Fig. 7**). These data suggest that 65°C is sufficient to damage molecules essential for growth that were stored in spores, although spores themselves have high heat resistance. We did not find time/temperature dependent effects of heat on outgrowth, except that an inverse correlation between the treatment and the burst-time was observed in the 80°C groups.

**Figure 7.**
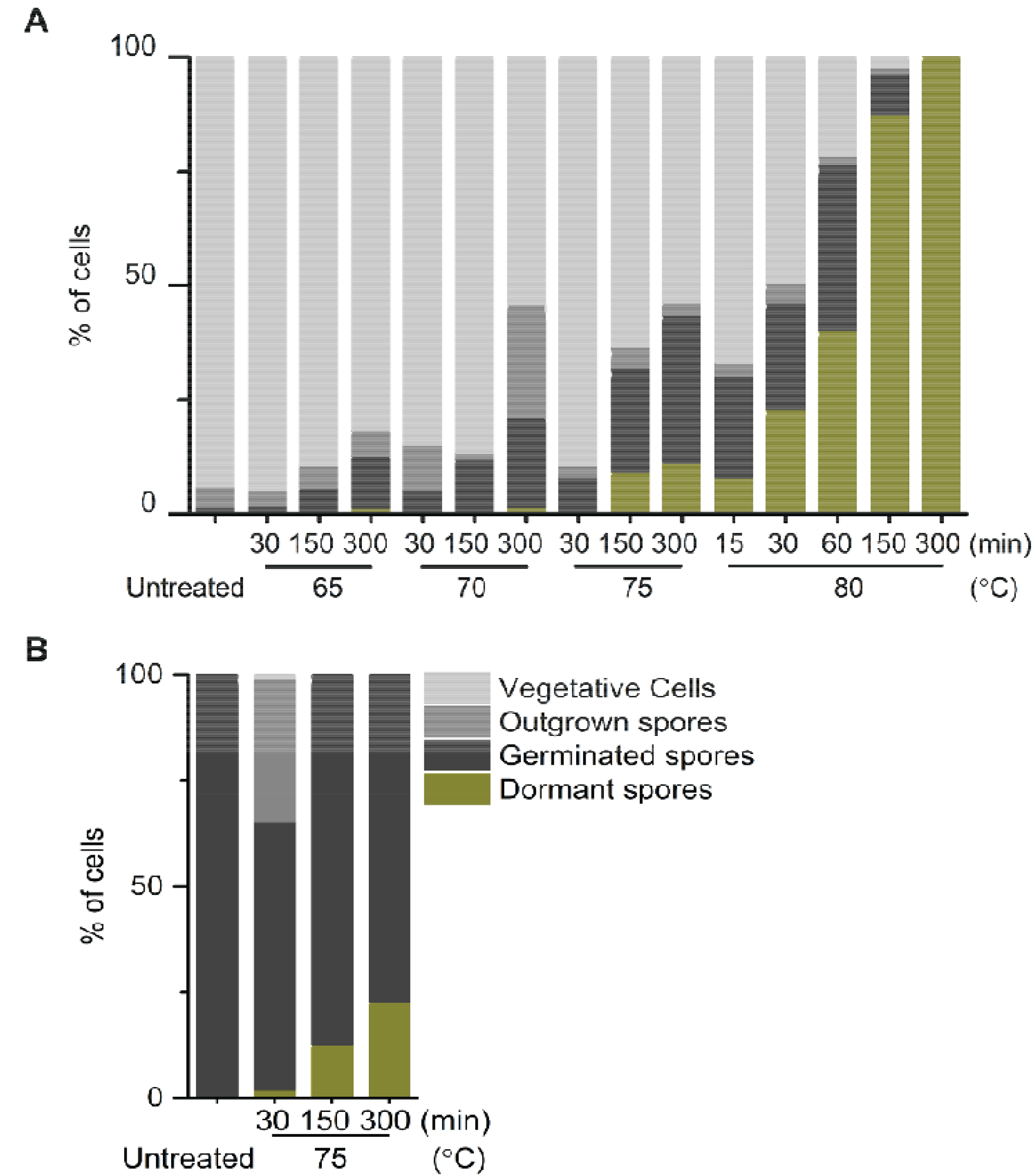
Quantification of spores with different fates after 16 hours of time lapse tracking. *B. subtilis* PS832 spores in water were heated at 65, 70, 75, or 80°C for different times. Subsequently, spores were immobilized on a MOPS-Agarose pad at 37°C with (10 mM each) AGFK (A) or 10 mM L-valine (B) for 16 hours of time lapse imaging. The number of individual spores examined by microscopy are given in Tables 1 and 2.

**Table 2.**
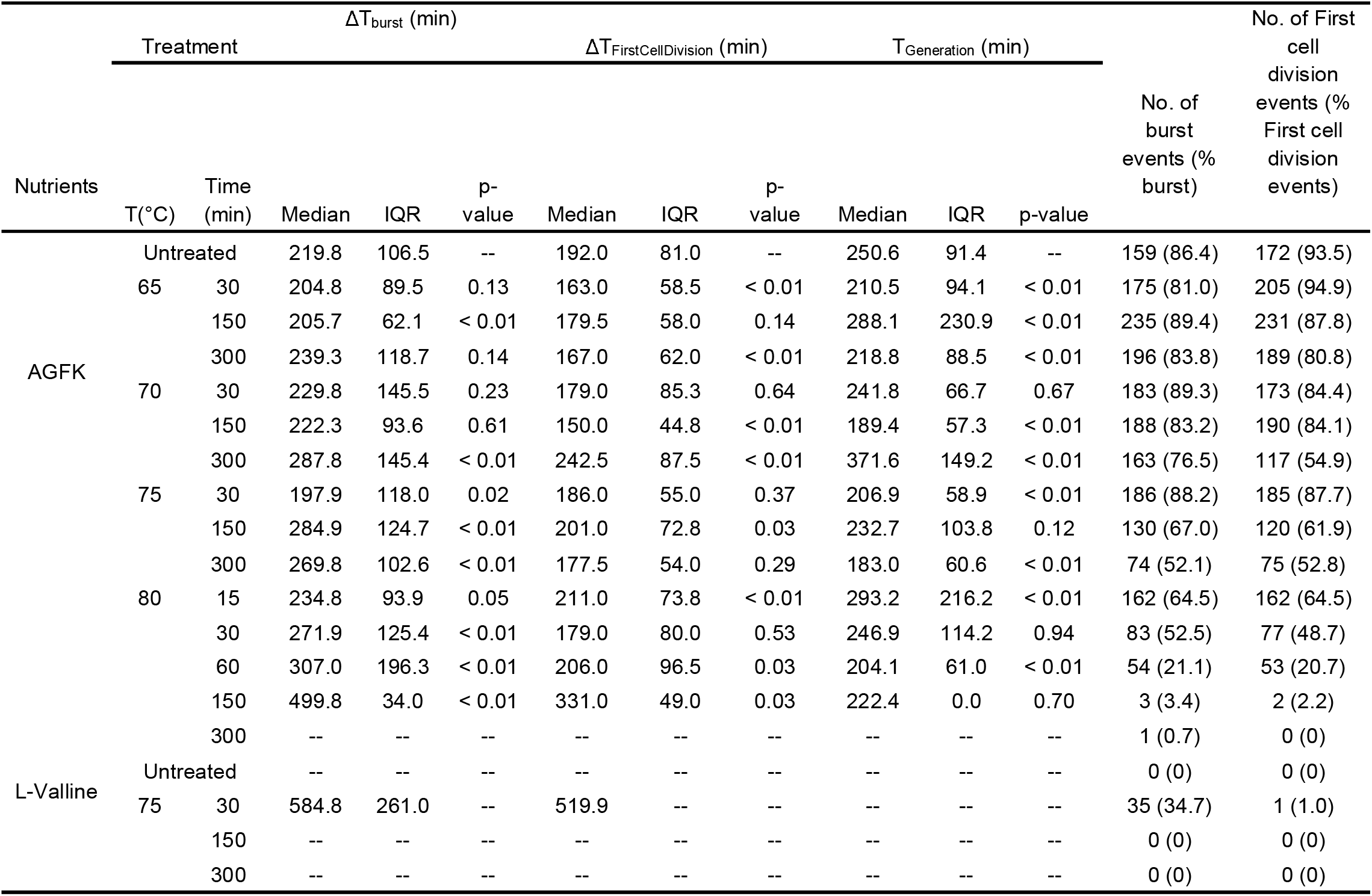
Outgrowth and growth parameters of individual heat treated *B. subtilis* PS832 spores in time lapse images^a^.

| Nutrients | Treatment | | $\Delta T_{burst}$ (min) | | | $\Delta T_{FirstCellDivision}$ (min) | | | $T_{Generation}$ (min) | | | No. of burst events (% burst) | No. of First cell division events (% First cell division events) |
| --- | --- | --- | --- | --- | --- | --- | --- | --- | --- | --- | --- | --- | --- |
|  |  |  | Median | IQR | p-value | Median | IQR | p-value | Median | IQR | p-value |  |  |
| AGFK | Untreated |  | 219.8 | 106.5 | -- | 192.0 | 81.0 | -- | 250.6 | 91.4 | -- | 159 (86.4) | 172 (93.5) |
|  | 65 | 30 | 204.8 | 89.5 | 0.13 | 163.0 | 58.5 | < 0.01 | 210.5 | 94.1 | < 0.01 | 175 (81.0) | 205 (94.9) |
|  |  | 150 | 205.7 | 62.1 | < 0.01 | 179.5 | 58.0 | 0.14 | 288.1 | 230.9 | < 0.01 | 235 (89.4) | 231 (87.8) |
|  |  | 300 | 239.3 | 118.7 | 0.14 | 167.0 | 62.0 | < 0.01 | 218.8 | 88.5 | < 0.01 | 196 (83.8) | 189 (80.8) |
|  | 70 | 30 | 229.8 | 145.5 | 0.23 | 179.0 | 85.3 | 0.64 | 241.8 | 66.7 | 0.67 | 183 (89.3) | 173 (84.4) |
|  |  | 150 | 222.3 | 93.6 | 0.61 | 150.0 | 44.8 | < 0.01 | 189.4 | 57.3 | < 0.01 | 188 (83.2) | 190 (84.1) |
|  |  | 300 | 287.8 | 145.4 | < 0.01 | 242.5 | 87.5 | < 0.01 | 371.6 | 149.2 | < 0.01 | 163 (76.5) | 117 (54.9) |
|  | 75 | 30 | 197.9 | 118.0 | 0.02 | 186.0 | 55.0 | 0.37 | 206.9 | 58.9 | < 0.01 | 186 (88.2) | 185 (87.7) |
|  |  | 150 | 284.9 | 124.7 | < 0.01 | 201.0 | 72.8 | 0.03 | 232.7 | 103.8 | 0.12 | 130 (67.0) | 120 (61.9) |
|  |  | 300 | 269.8 | 102.6 | < 0.01 | 177.5 | 54.0 | 0.29 | 183.0 | 60.6 | < 0.01 | 74 (52.1) | 75 (52.8) |
|  | 80 | 15 | 234.8 | 93.9 | 0.05 | 211.0 | 73.8 | < 0.01 | 293.2 | 216.2 | < 0.01 | 162 (64.5) | 162 (64.5) |
|  |  | 30 | 271.9 | 125.4 | < 0.01 | 179.0 | 80.0 | 0.53 | 246.9 | 114.2 | 0.94 | 83 (52.5) | 77 (48.7) |
|  |  | 60 | 307.0 | 196.3 | < 0.01 | 206.0 | 96.5 | 0.03 | 204.1 | 61.0 | < 0.01 | 54 (21.1) | 53 (20.7) |
|  |  | 150 | 499.8 | 34.0 | < 0.01 | 331.0 | 49.0 | 0.03 | 222.4 | 0.0 | 0.70 | 3 (3.4) | 2 (2.2) |
|  |  | 300 | -- | -- | -- | -- | -- | -- | -- | -- | -- | 1 (0.7) | 0 (0) |
| L-Valline | Untreated |  | -- | -- | -- | -- | -- | -- | -- | -- | -- | 0 (0) | 0 (0) |
|  | 75 | 30 | 584.8 | 261.0 | -- | 519.9 | -- | -- | -- | -- | -- | 35 (34.7) | 1 (1.0) |
|  |  | 150 | -- | -- | -- | -- | -- | -- | -- | -- | -- | 0 (0) | 0 (0) |
|  |  | 300 | -- | -- | -- | -- | -- | -- | -- | -- | -- | 0 (0) | 0 (0) |

#### 75°C treatment effects L-valine and AGFK induced germination in a similar way

At the population level, 75°C treatment promoted AGFK induced germination in a time dependent manner, which is different from results with L-valine triggered germination (**Fig. 1F, Fig. 2F**). To better understand this difference, L-valine induced germination was tracked by phase contrast microscopy after spore heating at 75°C. Unlike spores supplemented by AGFK and MOPS medium, no cell division events occurred in nutrient conditions with L-valine and MOPS. Similar to the cell growth in LB medium (**Fig. 4**), AGFK acted as nutritional assistance for cell growth. In both L-valine and AGFK induced germination, germination efficiency decreased with increased heat treatment time. In addition, T1 and ΔT_SlowLeakage_ were shortened in a time positive manner, and ΔT_PhaseDarkening_ was prolonged in a time positive manner (**Table 1**). In a word, the 75°C heat treatment affected AGFK and L-valine germination in the same way. The detectable time positive correlation in AGFK induced germination at population level was due to the bigger differences of germination parameters of different groups. However, it is not clear why AGFK induced germination is more sensitive to the change in heat activation time.

## Discussion

Heterogeneous spore germination in processed food is a major concern for food spoilage and foodborne disease. Promoting synchronous germination and then inactivating germinated spores and vegetative cells would extend food shelf life and at minimal cost in the decontamination process [10]. In the laboratory, spore germination heterogeneity can be stimulated by a heat activation procedure consisting of an exposure for a given time period to sublethal temperatures. However, with elevated temperatures, heat treatment could result in the accumulation of damage, and further increase in the germination and outgrowth heterogeneity [18,19]. Thus, in fact thermal treatment can create increased difficulties in food processing. In the current work, we measured the spore germination and growth profile after spore heat treatment at sublethal temperatures at a variety of treatment times. We hoped to determine the optimal heat time/temperature combination for optimal *B. subtilis* spore germination and outgrowth.

Our study showed that 50-65°C is the optimal heat activation temperature for *B. subtilis* PS832 spores obtained from cells sporulated in 2× SG medium at 37°C. The 50-65°C treatment resulted in the largest decrease in OD_595_ without any germination apparatus damage. Such damage, as well as damage to molecules essential for growth stored in the dormant spore, linked to increased levels of spore inactivation, starts accumulating after 70-75°C treatment. Published differential scanning calorimetry (DSC) thermogram profiles of *B. subtilis* spores present a reversible endothermic peak at around 60°C, which is followed by a second irreversible endothermic peak initiated at around 70°C [20]. It was suggested that the reversible endothermic peak is the outcome of heat activation, and the following irreversible peak is referred to as an inactivation peak. Our measured results fit the prediction from the *B. subtilis* thermogram profile. While we didn’t measure the heat activation/inactivation temperatures for other spore formers, it is likely that other *Bacillus* spores require similar heat activation/inactivation temperatures, based on their typical thermogram profile. Indeed, the activation endothermic peak for *Bacillus megaterium* spores is at 56°C, and for *Bacillus cereus* spores is at ∼55°C [21,22]. Notably, heat activation temperature has a positive correlation with heat resistance. For instance, super-dormant spores or very heat resistant spores require higher heat activation temperatures [5,23,24]. It would be useful to investigate the optimal heat activation conditions for heat resistant spores that are major troublemakers in the food industry [2].

Sublethal heat activation is reversible in a temperature dependent manner [25]. The following phenomena have been observed in heat activated spores, including but not limited to, some release of Ca^2+^DPA, a loss of coat proteins and small acid-soluble proteins, a change in the ultrastructure of the spore coat including the development of comblike striations on the coat, and reversible protein denaturation [13,26–30]. However, how heat activation works and why spores require 50-65°C to gain the optimal heat activation is still unclear. Studies have proposed heat activation might promote germination by directly increasing the level of functional GRs and affecting the state of the spore inner membrane, which surrounds the GRs [9,31]. Our data support the notion that GRs’ response to nutrient triggers has an important role in heat activation, considering the different germination behavior in L-valine and AGFK induced germination processes seen previously and in the current work.

The exact mechanism of spore heat activation remains unclear as is the full mechanism of heat damage to spores. One study showed that the inner membrane of spore was damaged in the spore heat inactivation process [32]. Other studies also suggested that moist heat (e.g. 89 °C for 2h) killed spores by protein denaturation in *B. subtilis, B. cereus, B. megaterium* and *Clostridium perfringens* spores [6,19,33]. Current work shows that effects of spores’ thermal treatment are different on different growth stages, including germination, outgrowth, and subsequent vegetative cell growth. Further study should focus on the relation between the thermal properties of bacterial spores and their proteome-wide scale analysis. This work might identify one or more macromolecules and/or processes key to the various thermal stress effects described in the current study.

## Materials and Methods

### Strain used and spore preparation

*B. subtilis* PS832, a prototrophic derivative of *B. subtilis* strain 168, was obtained from the laboratory of Prof. Peter Setlow. Spores were obtained from cells cultured at 37°C in 2× SG medium in Erlenmeyer flasks under continuous rotation at 200 rpm for 2 days using the procedure detailed by Abhyankar et al. [34]. Two day sporulation cultures were harvested, followed by extensive washing with MilliQ water, and further purification of phase bright spores using HistodenZ density gradient centrifugation [35]. Small volumes of aqueous spore suspensions (OD_600_ ∼ 60, in MIlliQ water) were dispensed, and frozen and stored at -80°C until use after confirming the phase brightness of the spores (≥ 98%) by phase contrast microscopy, as well as the absence of any visible debris.

### Heat treatment

The heat treatment procedure was modified based on Luu’s methods, in which spores were prepared at 37°C on 2× SG medium agar plates [9]. Aqueous spore suspensions (OD_600_ ∼ 2, ∼ 3×10^8^/ml) were incubated at 45, 50, 55, 60, 65, 70, 75 or 80 °C, for 0,15, 30, 60, 150, 240 and 300 min, followed by cooling in a water-ice bath (⍰15 min).

### Monitoring germination and growth at population level

The exchange of spore core Ca^2+^DPA for water during germination results in a drop of phase brightness, so that spore germination can be monitored by following spores’ optical density at 595 nm (OD_595_). Heat treated aqueous spore suspensions (final OD_595_ ∼ 1; 150 ⍰L) were dispensed into 96-well microtiter plates, and germinated (and grown) either in HEPES buffer (25 mM) or LB medium supplemented with either L-valine (10 mM) or AGFK (L-asparagine, glucose, fructose, and potassium chloride, 10 mM each). OD_595_ was measured every 5 min at 37°C from cultures that underwent continuous shaking in a Multiskan™ FC Microplate Photometer (Thermo Scientific). Data were collected from at least two independent experiments with at least two replicates for each individual experimental condition.

### Spore viability after heat treatment

Heat treated aqueous spore suspensions (OD_595_ ∼ 2) were diluted 1/10 in MilliQ water at room temperature. Aliquots of further dilutions were plated on LB agar plates which were incubated at 37°C overnight, followed by incubation at room temperature until no further colonies appeared. Colonies were counted to determine spore viability in least two independent biological experiments, with at least two replicates for each heat treatment condition.

### Monitoring germination, outgrowth and growth of individual spores

Heat treated PS832 spores (0.5 µL, OD_595_∼ 2) were immobilized on 1% agarose pads, supplemented with MOPS minimal medium and additional nutrient germinants (10 mM valine or 10 mM (each) AGFK), in an air containing chamber as described elsewhere [36]. Time-lapse phase contrast images were captured by a phase contrast microscope, which was coupled with a Nikon Ti microscope, a NA1.45 plan Apo *λ* 100× Oil Ph3 DM objective, a C11440□22CU Hamamatsu ORCA flash 4.0 camera, and the NIS elements software. Spores of each heat treatment condition were imaged twice at 37°C to track spore germination, outgrowth and growth in detail. For each imaging process, 6-9 fields of views were recorded in parallel once every 1 min for 16 h.

Images were analyzed by the modified ImageJ macro SporeTrackerX, which runs with the assistance of ImageJ plugin ObjectJ [13]. SporeTrackerX is capable of assessing multiple germination, outgrowth and growth events based on the drop of spore brightness, “jump-like” surface area increase, and cell surface area increase in phase contrast time-lapse images [13,36]. Briefly, the spore for analysis was labeled in the first frame of the time-lapse images, subsequently the changes of the brightness, as well as the log_2_(area) of the spore in time were assessed, stored, and plotted by SporeTrackerX, as shown in **Fig 1**. The conspicuous fast drop is almost certainly due to the rapid Ca^2+^DPA release and then subsequent cortex hydrolysis induced by the released Ca^2+^DPA (**Fig. 1A, C**) [4]. Both the time of the start of rapid Ca^2+^DPA release (T_StartRelease_) and the time of the appearance of the phase dark spore (T_PhaseDark_) were marked and stored for quantitative analysis. In addition, a slow decline in brightness occurred before the rapid drop, and the start time of the slow drop of brightness was stored as T_1_ (**Fig. 1A, C**). The slow drop is most likely due to the slow leakage Ca^2+^DPA [4]. In the growth plot, T_Burst_ was defined as the time of escape of the cell from the spore coat, T_FirstCellDivision_ was the time of the first cell division, and the generation time (T_Generation_) was the time between the first division until the end of the linear part of the log_2_(area) vs. time plot. (**Fig. B**). Based on the measurements described above, SporeTrackerX also calculated the following parameters: ⍰T_SlowLeakage_ is the time between T1 and T_StartRelease_.*⍰*T_PhaseDarkening_ is the time between T_StartRelease_ and T_PhaseDark_. *⍰T*_burst_ is the time between T_PhaseDark_ and T_Burst_.*⍰T*_FirstCellDivision_ is the time between T_Burst_ and T_FirstCellDivision_.

## Acknowledgements

We thank the Van Leeuwenhoek Centre for Advanced Microscopy at the University of Amsterdam for the use of the microscope. JW acknowledges the China Scholarship Council for a PhD fellowship.

## Author contributions

Juan Wen, funding acquisition. Juan Wen and Arend L. de Vos prepared the spore samples for investigation and formal analysis. Jan P. P. M. Smelt performed the modelling. Norbert O.E. Vischer created the image analysis software SporeTrackerX and assisted in the analyses. Juan Wen, Jan P. P. M. Smelt, and Arend L. De Vos wrote the original draft. Peter Setlow supervised the work revised and edited the manuscript. Stanley Brul, conceptualized, acquired funding and supervised the work, as well as wrote, reviewed and edited the manuscript.

## References

1. Smelt Jppm, JPP, Brul S. Thermal inactivation of microorganisms. Crit Rev Food Sci Nutr. 2014;50: 1371–1385.

2. Wells-Bennik MHJ, Eijlander RT, den Besten HMW, Berendsen EM, Warda AK, Krawczyk AO, et al. Bacterial spores in food: survival, emergence, and outgrowth. Annu Rev Food Sci Technol. 2016;7: 457–482.

3. Abhyankar W, Pandey R, Ter Beek A, Brul S, de Koning LJ, de Koster CG. Reinforcement of Bacillus subtilis spores by cross-linking of outer coat proteins during maturation. Food Microbiol. 2015;45: 54–62.

4. Setlow P, Wang S, Li Y-Q. Germination of spores of the orders Bacillales and Clostridiales. Annu Rev Microbiol. 2017;71: 459–477.

5. Rodriguez-Palacios A, Lejeune JT. Moist-heat resistance, spore aging, and superdormancy in Clostridium difficile. Appl Environ Microbiol. 2011;77: 3085–3091.

6. Smelt Jppm, JPP, Bos AP, Kort R, Brul S. Modelling the effect of sub(lethal) heat treatment of Bacillus subtilis spores on germination rate and outgrowth to exponentially growing vegetative cells. Int J Food Microbiol. 2008;128: 34–40.

7. Levinson HS, Hyatt MT. Effects of temperature on activation, germination, and outgrowth of Bacillus megaterium spores. J Bacteriol. 1970;101: 58–64.

8. Keynan A, Evenchik Z, Halvorson HO, Hastings JW. Activation of bacterial endospores. J Bacteriol. 1964;88: 313–318.

9. Luu S, Cruz-Mora J, Setlow B, Feeherry FE, Doona CJ, Setlow P. The effects of heat activation on Bacillus spore germination, with nutrients or under high pressure, with or without various germination proteins. Appl Environ Microbiol. 2015;81: 2927–2938.

10. Løvdal IS, Hovda MB, Granum PE, Rosnes JT. Promoting Bacillus cereus spore germination for subsequent inactivation by mild heat treatment. J Food Prot. 2011;74: 2079–2089.

11. Paidhungat M, Setlow P. Role of ger proteins in nutrient and nonnutrient triggering of spore germination in Bacillus subtilis. J Bacteriol. 2000;182: 2513–2519.

12. Black EP, Koziol-Dube K, Guan D, Wei J, Setlow B, Cortezzo DE, et al. Factors influencing germination of Bacillus subtilis spores via activation of nutrient receptors by high pressure. Appl Environ Microbiol. 2005;71: 5879–5887.

13. Omardien S, Ter Beek A, Vischer N, Montijn R, Schuren F, Brul S. Evaluating novel synthetic compounds active against Bacillus subtilis and Bacillus cereus spores using Live imaging with SporeTrackerX. Sci Rep. 2018;8: 1–13.

14. Leuschner RGK, Lillford PJ. Effects of temperature and heat activation on germination of individual spores of Bacillus subtilis. Lett Appl Microbio. 1999;29: 228–232.

15. Sinai L, Rosenberg A, Smith Y, Segev E, Ben-Yehuda S. The molecular timeline of a reviving bacterial spore. Mol Cell. 2015;57: 695–707.

16. Zhang P, Thomas S, Li Y-Q, Setlow P. Effects of cortex peptidoglycan structure and cortex hydrolysis on the kinetics of Ca2 -Dipicolinic acid release during Bacillus subtilis spore germination. J Bacteriol. 2012;194: 646–652.

17. Wang S, Setlow P, Li Y-Q. Slow leakage of Ca-dipicolinic acid from individual Bacillus spores during initiation of spore germination. J Bacteriol. 2015;197: 1095–1103.

18. Warda AK, den Besten HMW, Sha N, Abee T, Nierop Groot MN. Influence of food matrix on outgrowth heterogeneity of heat damaged Bacillus cereus spores. Int J Food Microbiol. 2015;201: 27–34.

19. He L, Chen Z, Wang S, Wu M, Setlow P, Li Y-Q. Germination, outgrowth, and vegetative growth kinetics of dry heat treated Individual Spores of Bacillus Species. Appl Environ Microbiol. 2018;84.

20. Ablett S, Darke AH, Lillford PJ, Martin DR. Glass formation and dormancy in bacterial spores. Int J Food Sci. 1999;34: 59–69.

21. Leuschner RGK, Lillford PJ. Thermal properties of bacterial spores and biopolymers. Int J Food Microbiol. 2003;80: 131–143.

22. Belliveau BH, Beaman TC, Pankratz HS, Gerhardt P. Heat killing of bacterial spores analyzed by differential scanning calorimetry. J Bacteriol. 1992;174: 4463–4474.

23. Berendsen EM, Zwietering MH, Kuipers OP, Wells-Bennik MHJ. Two distinct groups within the Bacillus subtilis group display significantly different spore heat resistance properties. Food Microbiol. 2015;45: 18–25.

24. Ghosh S, Zhang P, Li Y-Q, Setlow P. Superdormant spores of Bacillus species have elevated wet-heat resistance and temperature requirements for heat activation. J Bacteriol. 2009;191: 5584–5591.

25. Busta FF, John Ordal Z. Heat-Activation Kinetics of Endospores of Bacillus subtilis. J Food Sci. 1964;29: 345–353.

26. Alimova A, Katz A, Gottlieb P, Alfano RR. Proteins and dipicolinic acid released during heat shock activation of Bacillus subtilis spores probed by optical spectroscopy. Appl Opt. 2006;45: 445–450.

27. Zhang P, Setlow P, Li Y. Characterization of single heat-activated Bacillus spores using laser tweezers Raman spectroscopy. Opt Express. 2009;17: 16480–16491.

28. Wang G, Zhang P, Paredes-Sabja D, Green C, Setlow P, Sarker MR, et al. Analysis of the germination of individual Clostridium perfringens spores and its heterogeneity. J Appl Microbiol. 2011;111: 1212–1223.

29. Zhang P, Kong L, Setlow P, Li Y-Q. Characterization of wet-heat inactivation of single spores of bacillus species by dual-trap Raman spectroscopy and elastic light scattering. Appl Environ Microbiol. 2010;76: 1796–1805.

30. Luu S, Setlow P. Analysis of the loss in heat and acid resistance during germination of spores of Bacillus species. J Bacteriol. 2014;196: 1733–1740.

31. Zhang P, Garner W, Yi X, Yu J, Li Y-Q, Setlow P. Factors affecting variability in time between addition of nutrient germinants and rapid dipicolinic acid release during germination of spores of Bacillus species. J Bacteriol. 2010;192: 3608–3619.

32. Fan L, Ismail BB, Hou F, Muhammad AI, Zou M, Ding T, et al. Thermosonication damages the inner membrane of Bacillus subtilis spores and impels their inactivation. Food Res Int. 2019;125: 108514.

33. Wang G, Paredes-Sabja D, Sarker MR, Green C, Setlow P, Li Y-Q. Effects of wet heat treatment on the germination of individual spores of Clostridium perfringens. J Appl Microbiol. 2012;113: 824–836.

34. Abhyankar WR, Kamphorst K, Swarge BN, van Veen H, van der Wel NN, Brul S, et al. The influence of sporulation conditions on the spore coat protein composition of Bacillus subtilis spores. Frontiers in Microbiology. 2016.01636

35. Wen J, Pasman R, Manders EMM, Setlow P, Brul S. Visualization of germinosomes and the inner membrane in Bacillus subtilis spores. JoVE Journal Vis. Exp. 2019. e59388.

36. Pandey R, Ter Beek A, Vischer NOE, Jan P P, Brul S, Manders EMM. Live cell imaging of germination and outgrowth of individual Bacillus subtilis spores; the effect of heat stress quantitatively analyzed with SporeTracker. PLoS One. 2013;8: e58972.

